# Quantification of frequency-dependent genetic architectures and action of negative selection in 25 UK Biobank traits

**DOI:** 10.1101/188086

**Authors:** Armin P Schoech, Daniel Jordan, Po-Ru Loh, Steven Gazal, Luke O’Connor, Daniel J Balick, Pier F Palamara, Hilary K Finucane, Shamil R Sunyaev, Alkes L Price

**Affiliations:** Department of Epidemiology, Harvard T.H. Chan School of Public Health, Boston, Massachusetts, USA; Program in Medical and Population Genetics, Broad Institute of Harvard and MIT, Cambridge, Massachusetts, USA; Department of Genetics and Genomic Sciences, Icahn School of Medicine at Mount Sinai, New York City, New York, USA; Division of Genetics, Department of Medicine, Brigham and Women’s Hospital, Harvard Medical School, Boston, Massachusetts, USA; Department of Statistics, University of Oxford, Oxford, United Kingdom

## Abstract

Understanding the role of rare variants is important in elucidating the genetic basis of human diseases and complex traits. It is widely believed that negative selection can cause rare variants to have larger per-allele effect sizes than common variants. Here, we develop a method to estimate the minor allele frequency (MAF) dependence of SNP effect sizes. We use a model in which per-allele effect sizes have variance proportional to [*p*(1−*p*)]^α^, where *p* is the MAF and negative values of *α* imply larger effect sizes for rare variants. We estimate *α* by maximizing its profile likelihood in a linear mixed model framework using imputed genotypes, including rare variants (MAF >0.07%). We applied this method to 25 UK Biobank diseases and complex traits (N = 113,851). All traits produced negative *α* estimates with 20 significantly negative, implying larger rare variant effect sizes. The inferred best-fit distribution of true *α* values across traits had mean −0.38 (s.e. 0.02) and standard deviation 0.08 (s.e. 0.03), with statistically significant heterogeneity across traits (P = 0.0014). Despite larger rare variant effect sizes, we show that for most traits analyzed, rare variants (MAF <1%) explain less than 10% of total SNP-heritability. Using evolutionary modeling and forward simulations, we validated the *α* model of MAF-dependent trait effects and estimated the level of coupling between fitness effects and trait effects. Based on this analysis an average genome-wide negative selection coefficient on the order of 10^−4^ or stronger is necessary to explain the *α* values that we inferred.

## Introduction

The contribution of rare variants to the genetic architecture of human diseases and complex traits is a question of fundamental interest, which can inform the design of genetic association studies and shed light on the action of negative selection^1, 2^. Recently, several studies have investigated the relationship between minor allele frequency (MAF) and trait effects^3, 4, 5, 6^. However, these studies have analyzed a small number of traits and have not evaluated the genome-wide contribution of rare variants (MAF < 1%), which remains unknown^7^.

Here we develop a profile likelihood-based mixed model method to infer MAF-dependent architectures from genotype and phenotype data. We apply our method to 25 complex traits and diseases from the UK Biobank data set, analyzing data from 113,851 individuals and 11,062,620 SNPs, including rare variants (MAF > 0.07%). Our analysis shows that rare variants have significantly increased per-allele effect sizes for most traits, with significant heterogeneity across traits. For each of these traits we also estimate the phenotypic variance explained by variants in different frequency ranges, including rare variants.

It is widely believed that frequency-dependence of SNP effect sizes is due to increased negative selection on variants that affect complex traits^1, 2, 8, 9, 10, 11^. Specifically, if SNPs that affect a trait are more likely to be under negative selection, they will be enriched in the lower-frequency spectrum, so that lower-frequency SNPs will on average have larger trait effects. Thus, MAF-dependent architectures estimated from genotype and phenotype data can shed light on evolutionary parameters. Previous studies have used MAF-dependent architectures or related information to estimate a coupling parameter^9^ between fitness effects and their trait effects for prostate cancer^6^ and type 2 diabetes^12, 13^. In this work, we use evolutionary modeling and forward simulations to investigate whether our parameterization of MAF-dependent effects (α model; see below) is consistent with evolutionary models, estimate the coupling between fitness effects and trait effects, and draw inferences about the average genome-wide strength of negative selection.

## Results

### Overview of methods

We assume a previously proposed random-effect model^14, 15^ (the “α model”), in which the per-allele trait effect β of a SNP depends on its MAF *p* via:

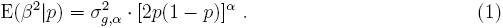

A negative value of α implies that lower-frequency SNPs have larger per-allele effect sizes, whereas α = 0 implies no dependence, and 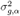 is the component of SNP effect variance that is independent of frequency. We note that Equation 1 pertains to genome-wide SNPs, including SNPs that do not affect the trait. The α model is simple and convenient, but has not previously been validated by evolutionary modeling.

For a given set of genotype and phenotype data, we estimate α using a linear mixed model framework^16^. The model likelihood depends on α, 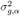, and the environmental variance (see Online Methods). We compute the profile likelihood over values of α by maximizing the likelihood with respect to and the environmental variance for a given α. Our estimate 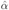 is defined as the mode of the profile likelihood curve, whose width is used to compute error estimates. We show that the corresponding values of 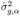 can be used to estimate the SNP-heritability 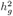 while accounting for MAF-dependent SNP effects, which can bias 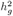 estimates when not accounted for^14, 15^. We include linkage disequilibrium (LD)-dependent SNP weights^17^ in our model, to avoid biases due to LD-dependent architectures^4, 14, 18, 19^. Details of the method are described in the Online Methods section; we have released open-source software implementing the method (see URLs).

### Simulations

We evaluated our method using simulations based on imputed UK Biobank genotypes^20^ and simulated phenotypes, using *N* = 5,000 individuals and *M* = 100,000 consecutive SNPs from a 25Mb block of chromosome 1 (see Online Methods). We used default parameter settings of α = −0.3, 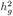 = 0.4, 1% of SNPs causal, imputation noise based on actual imputed genotype probabilities, and LD-dependent effects^17^, but we also considered other parameter settings for each of these. Imputation noise was introduced by randomly sampling the genotypes used to simulate phenotypes from imputed genotype probabilities, while still using the expected dosage values for inference (see Online Methods).

In Table 1, we report α estimates at default and other parameter settings, both using LD-dependent weights (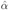) and without using LD-dependent weights (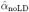). In simulations with LD-dependent effects, 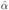 was unbiased at all parameter settings tested, while 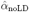 was upward biased by approximately 0.1. In simulations without LD-dependent effects, 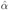 was downward biased by less than 0.1, while 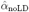 was unbiased. These simulations suggest that our method provides unbiased estimates of α when LD is correctly modeled, and only modestly biased estimates of α when LD is not correctly modeled. We also compared our profile likelihood standard error estimates to empirical standard errors from simulations. These quantities do not differ significantly (see Supplementary Table 1), indicating that our standard error estimates are well-calibrated. The profile likelihood curves were smooth and unimodal at all parameter settings (see Supplementary Figure 1).

**Table 1:**
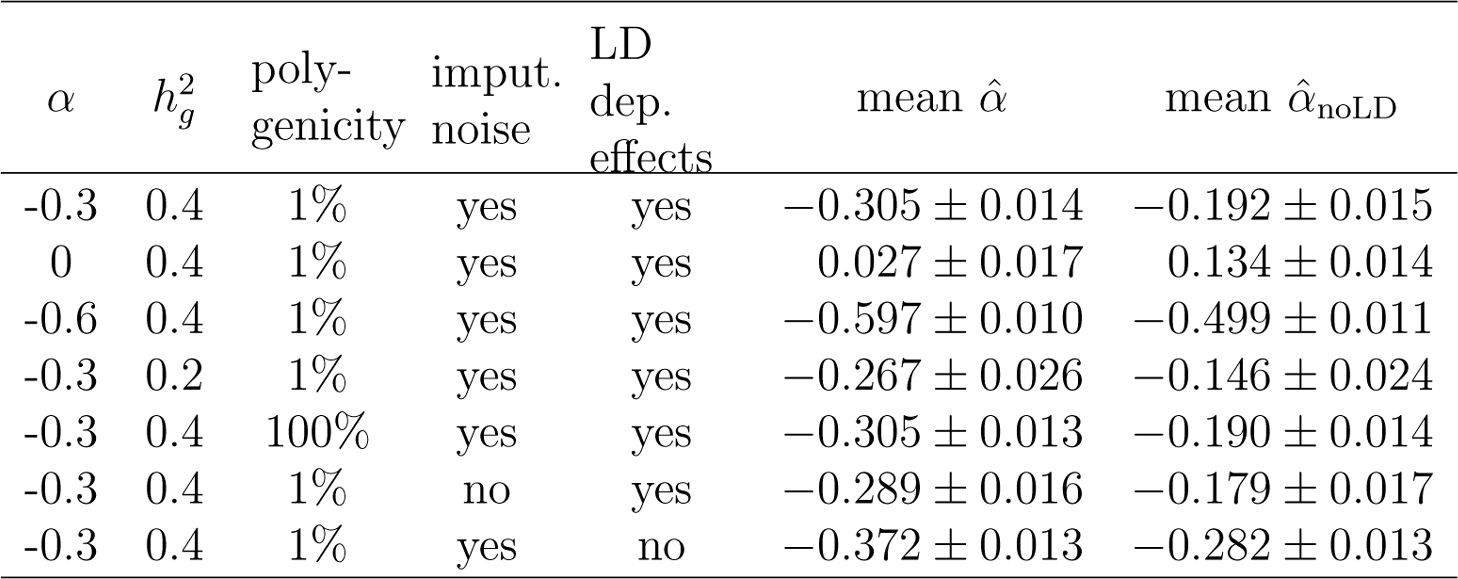
Estimates of α in simulations We simulated phenotypes using imputed UK Biobank genotypes and applied our method to infer α. In each line we show results from phenotypes that were simulated using various values of α, 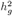, and the proportion of causal SNPs. In most simulations, imputation noise and LD dependent SNP effects were included in the simulated phenotypes. In each case we report the mean estimated α and standard error of the mean, using our estimation method either with LD correction (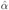) or without LD correction (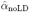).

Although the main focus of this paper is on obtaining and interpreting estimates of α, we also used our simulation framework to evaluate the effectiveness of our method in obtaining SNP-heritability estimates that avoid biases due to MAF-dependent and LD-dependent architectures. In Supplementary Table 2 we report SNP-heritability estimates using our method, both using LD-dependent weights (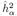) and without using LD-dependent weights (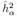), and using GCTA with a single variance component (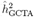)^16^. 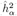 and 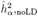 were roughly unbiased at all parameter settings, while GCTA with a single variance component produced biased estimates, consistent with previous work^4, 14^. Other methods of avoiding bias due to MAF-dependent and LD-dependent architectures have recently been proposed, including GREML-LDMS^4^ and LDAK^19^; a complete benchmarking of SNP-heritability estimation methods will be provided elsewhere (ref.^21^, which we are updating to include a comparison to LDAK^19^).

### Analysis of 25 UK Biobank traits

We applied our method to 113,851 British-ancestry individuals from the UK Biobank with 1000 Genomes- and UK10K-imputed genotypes at 11,062,620 SNPs with at least 5 minor alleles in the UK10K reference panel (MAF > 0.07%; see Online Methods). We analyzed 25 heritable, polygenic traits with at least 50% of individuals phenotyped (Table 2). Phenotype values were corrected for fixed effects, including sex and 10 principal components (see Online Methods). Profile likelihood curves for all 25 traits are displayed in Supplementary Figure 2. We observed that the curves were smooth and unimodal (consistent with simulations; Supplementary Figure 1), suggesting that estimates of α are likely to be robust.

**Table 2:**
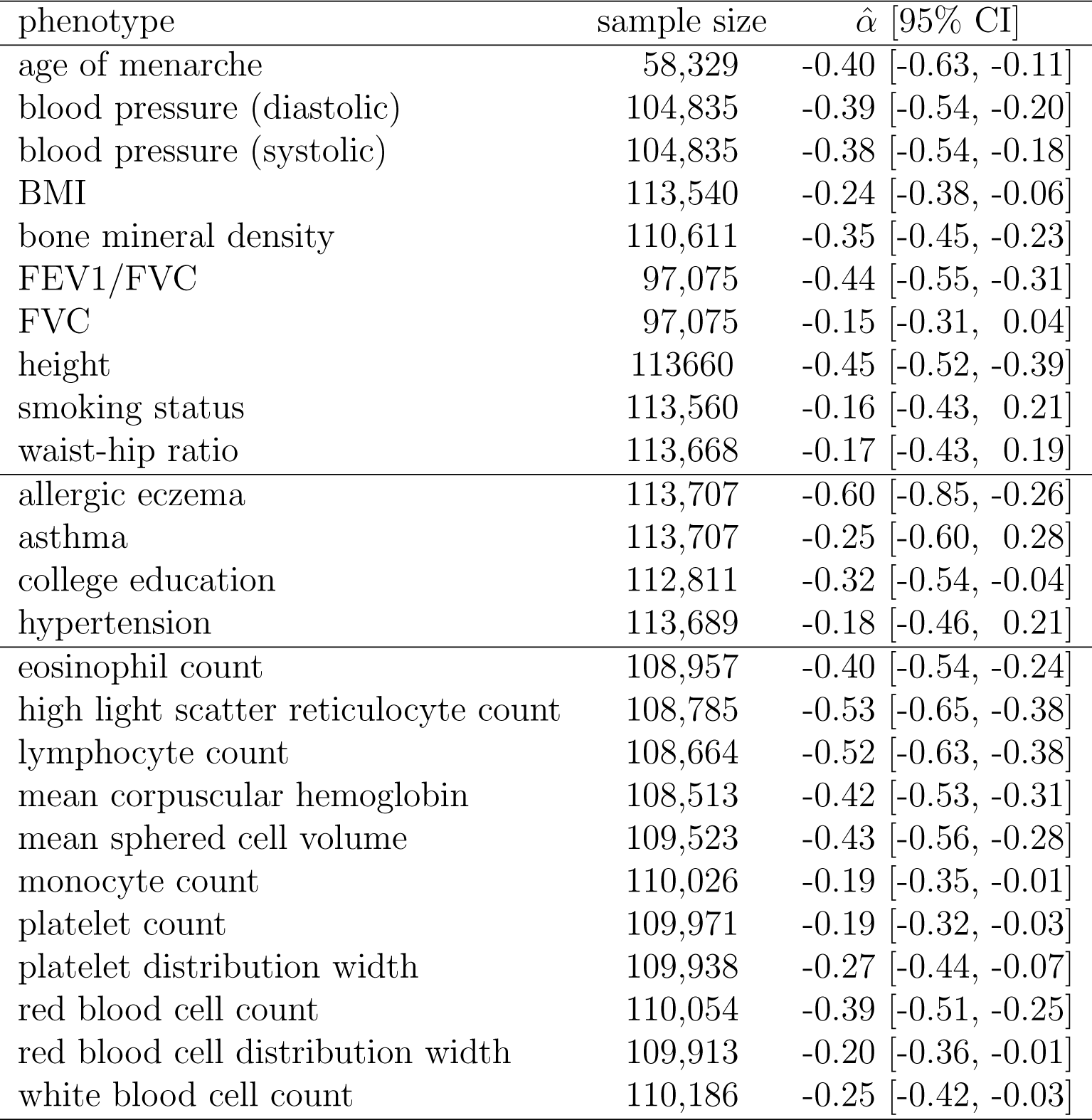
Estimates of α for 25 UK Biobank traits We computed α estimates for 25 UK Biobank traits, including 10 quantitative traits, 4 case-control traits, and 11 blood cell traits (all quantitative). The reported 95% credible intervals were calculated from the profile likelihood curves using a at prior.

In Table 2, we report estimates of α for all 25 traits. All traits had negative α estimates (with most estimates lying between −0.5 and −0.2), and 20 traits had significantly negative estimates (i.e. 95% credible intervals did not overlap zero), implying that lower-frequency SNPs have larger per-allele effect sizes. We observed statistically significant heterogeneity in estimates of α across the 25 traits (P = 0.0014), consistent with different levels of (direct and/or pleiotropic) negative selection across traits (see Discussion). We estimated the underlying distribution of true (unobserved) values of α to have mean −0.38 (s.e. 0.02) and standard deviation 0.08 (s.e. 0.03), assuming a normal distribution (see Online Methods). We obtained very similar results when repeating the entire analysis using 9,336,687 SNPs with MAF > 0.3% in the UK10K reference panel (Supplementary Table 3); we note that these results are unlikely to be affected by imputation error, because simulation results in Table 1 show that our method is not significantly affected by imputation error under correctly calibrated imputation accuracies, and because we further determined that MAF > 0.3% SNPs generally have well-calibrated imputation accuracies (Supplementary Figure 3).

We estimated the proportion of SNP-heritability explained by SNPs in each part of the MAF spectrum, for different values of α. This computation relies on the empirical MAF spectrum in UK10K, as heritability per MAF bin depends both on heritability per SNP and number of SNPs per MAF bin (see Online Methods). Results are reported in Figure 1. We determined that rare and low-frequency variants contribute a very small proportion of SNP-heritability at the mean α estimate of −0.38, and a relatively small proportion of SNP-heritability even for the most negative α estimate of −0.60. Specifically, at α = −0.38 (s.d. 0.08), only 8.9% (s.d. 2.7%) of SNP-heritability is explained by SNPs with MAF < 1%. We also used 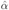 to obtain total SNP-heritability estimates corrected for biases due to MAF-dependent and LD-dependent architectures for each of the 25 traits (Supplementary Table 4; see Online Methods).

**Figure 1:**
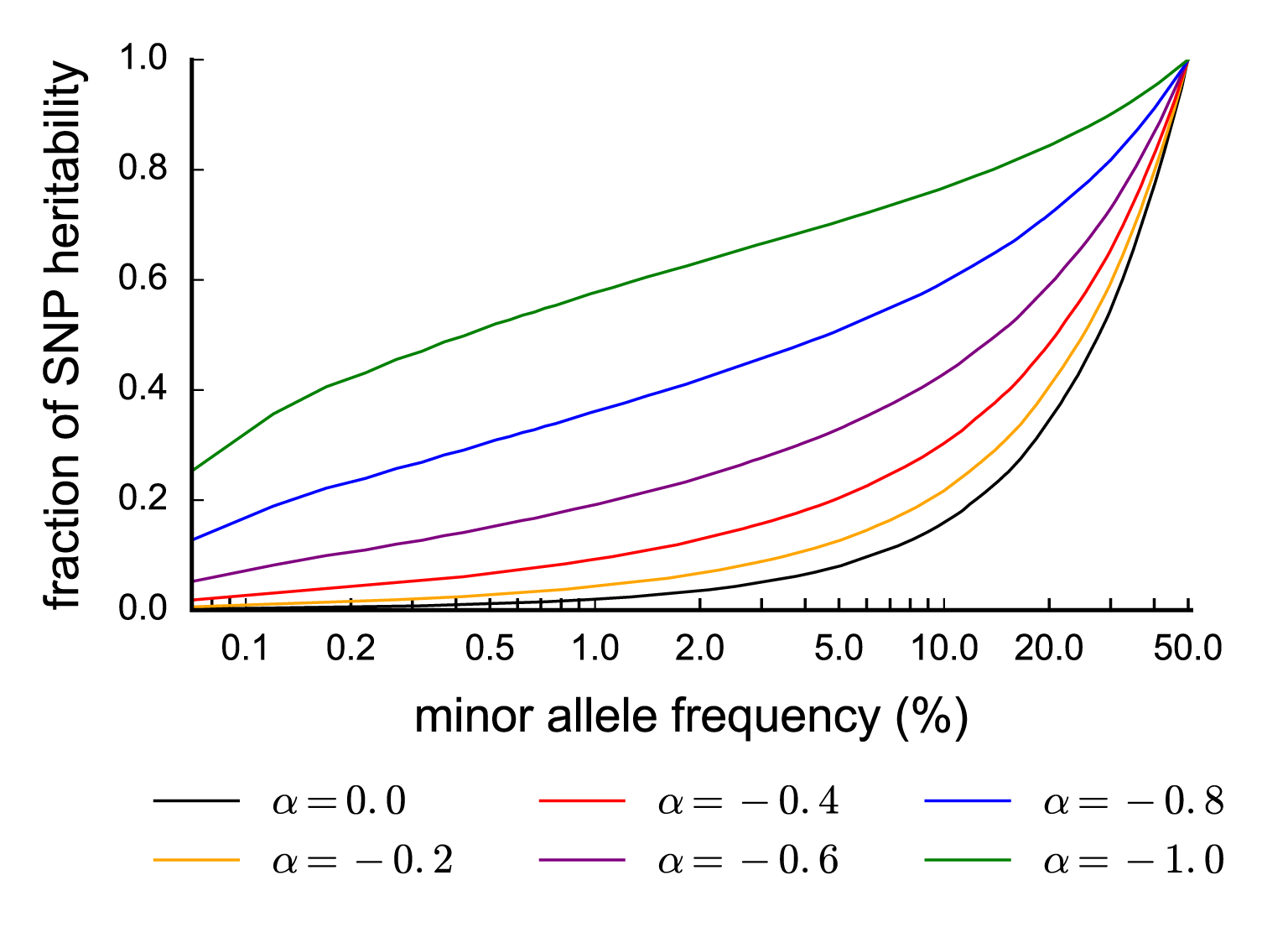
Fraction of SNP-heritability in different MAF ranges given α. We report the fraction of SNP-heritability explained by SNPs up to a certain MAF (x-axis), for different values of α. For example, assuming α = −0.4, SNPs with MAF ≤ 5% collectively explain about 20% of the total SNP-heritability. These results are based on the UK10K allele frequency spectrum and our model assumption that the squared per-allele effects is proportional to [2*p*(1 − *p*)]^α^.

### Effect of negative selection on the MAF-dependence of genetic architectures

Frequency-dependent trait effect sizes have been widely attributed to negative (purifying) selection on variants that affect complex traits, which causes them to be enriched for lower-frequency variants, so that lower-frequency SNPs will have larger traits effects^1, 2, 8, 9, 10, 11^. Here we use evolutionary modeling to predict the frequency-dependent architecture of a trait, given the coupling between fitness effects and trait effects. The aim of this analysis was to investigate whether the α model (Equation 1) is consistent with the predictions of evolutionary models, and to draw conclusions about evolutionary parameters from our estimates of α across 25 UK Biobank traits.

We used an evolutionary model of Eyre-Walker^9^, which introduces a parameter τ quantifying the coupling between a SNP’s fitness effect (selection coefficient *s*) and target trait effect size (β); τ > 0 implies that SNPs under negative selection have larger trait effect sizes on average, whereas τ = 0 corresponds to no coupling. Using this model, we derived two analytical results. First, it is straightforward to show that

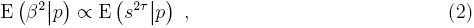

where *p* is minor allele frequency (see Online Methods). This implies that increased trait effects for lower-frequency variants requires both that lower-frequency variants have significantly larger selection coefficients s and that τ > 0. Second, based on Equation 2, we analytically evaluated E(*s*^2τ^|*p*) to quantify the MAF-dependence of SNP effects under the Eyre-Walker model (see Online Methods). In this derivation, we ignored LD between selected SNPs, assumed a constant effective population size *N_e_*, and assumed that selection coefficients *s* of SNP loci across the genome are drawn from a gamma distribution, with mean 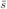 and shape parameter κ (ref.^22^). (We note that κ parametrizes the polygenicity of fitness: if κ » 1, all SNPs in the genome have roughly the same selection coefficient; if κ « 1, a few SNPs have extremely large selection coefficients.) Under these assumptions, we derived the result that there exists a MAF threshold *T* such that for *p* > *T* the α model approximately holds, but for *p* < *T* trait effects are approximately independent of frequency (see Online Methods). The threshold is

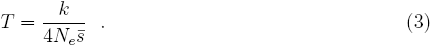

Intuitively, this threshold corresponds to the maximum frequency at which even the most strongly selected SNPs are still only affected by genetic drift, with their frequency being too low to be significantly affected by selection. We note that *T* is independent of the trait analyzed, since 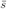 and κ parametrize the distribution of genome-wide selection coefficients.

Although our derivation of Equation 3 ignored the effects of demographic changes and LD, we confirmed this result by performing forward simulations using SLiM2^23^, using a European demographic model^24^ and realistic LD patterns (see Online Methods). Specifically, for a given τ we computed E (*s*^2τ^|*p*) in Equation 2 from the *s* and *p* values of simulated SNPs. Our main simulations assumed τ = 0.4, effective *N_e_* = 10,000, 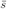 = 0.001 and κ = 0.25 (ref.^22^), so that *T* = 0.006 (Equation 3). Results are reported in Figure 2, which shows that for *p* > *T* = 0.006 the α model with best-fit α = −0.32 provides a good fit, but for *p* < *T* = 0.006 the effect sizes are less MAF-dependent and are thus significantly smaller than expected under the α model. Results at other parameter settings were qualitatively similar, with the threshold varying according to Equation 3 (see Supplementary Figure 4).

**Figure 2:**
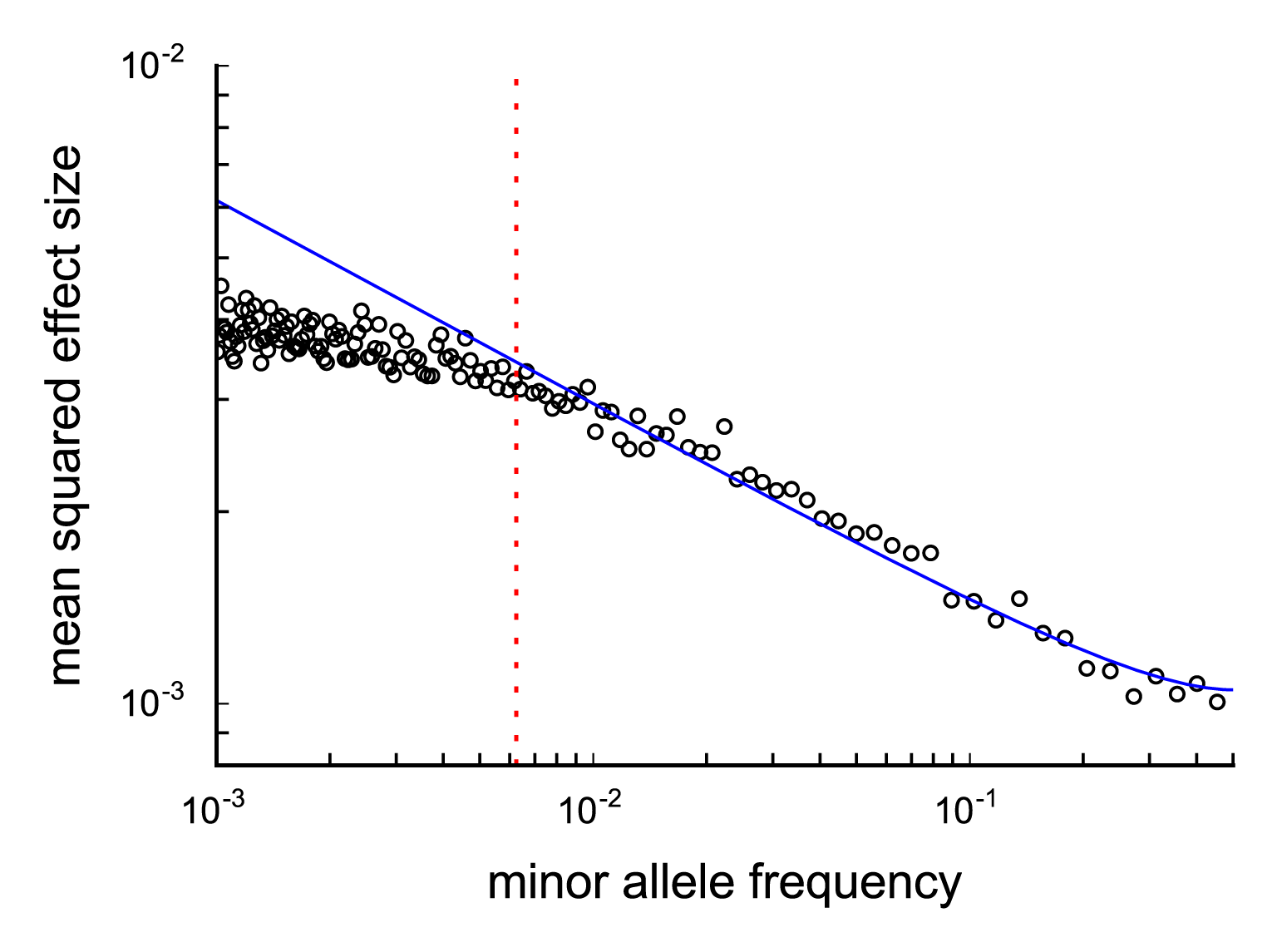
MAF-dependence of SNP effects in evolutionary forward simulations. Forward simulations confirm that α model approximately holds above the MAF threshold 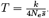. We report simulated mean squared SNP effect sizes at a given MAF on a log-log plot, assuming τ = 0.4 and a genome wide selection coefficient distribution with mean 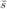 = 10^−3^ and shape parameter τ = 0.25. Data points represent the mean squared effect size of 1000 SNPs of similar MAF, calculated assuming Equation 2. The blue curve represents mean squared effect sizes under the α model (Equation 1) with α = −0.32, fitted to SNPs above the MAF threshold *T*. The MAF threshold *T* = 0.006 is indicated by a dotted red line.

We sought to draw inferences about the threshold *T* from our analysis of 25 traits. If a significant fraction of SNPs used to estimate α in that analysis had MAF below *T*, we would expect to obtain smaller (more negative) estimates of α by restricting to more common SNPs, since SNPs of MAF below *T* with less MAF-dependent effects would be ignored. We repeated the estimation of α for all 25 traits using 6,273,557 SNPs with MAF > 5% (instead of 11,062,620 SNPs with MAF > 0.07%). Estimates did not significantly change for any trait (see Supplementary Table 5), nor did the best-fit α estimate across traits, which actually increased slightly from −0.38 (s.e. 0.02) to −0.35 (s.e. 0.02). It is possible that effects of rare SNPs above the original MAF threshold of 0.07% are indeed overestimated in the α model (if *T* > 0.07%), but if so the impact of this deviation is not large enough to significantly change our estimates. On the other hand, it is unlikely that this is the case for all rare and low-frequency SNPs (MAF < 5%), since they explain roughly 10% of heritability even under a neutral model (Figure 1). We conclude that the threshold *T* is likely to be < 5%, so that the α model provides a good fit for common SNPs (MAF ≥ 5%). However, the α model may potentially overestimate the effects of rare SNPs. This implies that the fraction of heritability explained by rare SNPs in Figure 1 should be viewed as an upper bound.

Finally, we sought to draw conclusions about the values of the average genome-wide selection coefficient 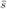 and the Eyre-Walker coupling parameter τ. First, a threshold *T* < 5% (see above) implies an average selection coefficient 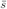 > 5κ/*N_e_*. Assuming *N_e_* = 10,000 (ref.^25^) and κ = 0.25 (ref.^22^), 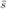 is likely to be on the order of 10^−4^ or stronger. Second, we determined that the best-fit estimate of 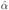 = −0.38 across 25 traits corresponds to a τ value in the range [0.3,0.5] (Figure 3, see Online Methods). We reached this conclusion by repeating our forward simulations for τ ∈ [0,1] (vs. τ = 0.4 above), 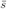 ∈ {0.0001, 0.001} (vs. 0.001 above) and τ ∈ {0.125, 0.25} (vs. 0.25 above) and fitting the α model using SNPs above the threshold *T* from Equation 3. Figure 3 shows that the best-fit α depends primarily on τ, with only weak dependence on 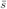 and κ. Estimates of τ for each of the 25 traits are provided in Supplementary Table 6.

**Figure 3:**
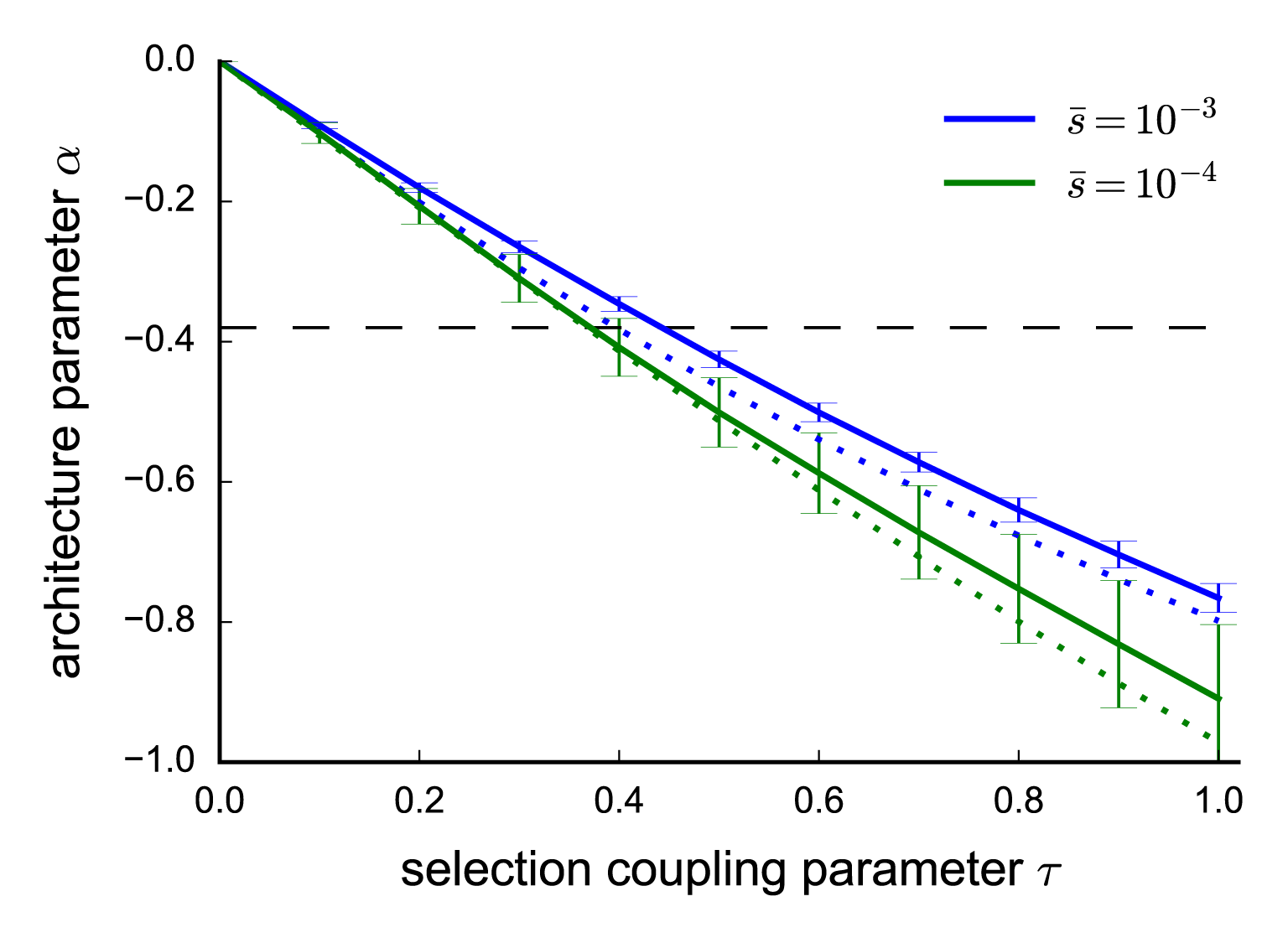
Value of α as a function of τ and other parameters in forward simulations. We report best-fit α estimates for simulations at each value of τ at a given genome-wide average selection coefficient 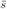. Selection coefficients were sampled using a gamma distribution shape parameter of τ = 0.25 (solid lines) or τ = 0.125 (dotted lines). α estimates where calculated by fitting the model in Equation 1 to simulated SNP effects above the MAF threshold 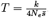, with error bars representing standard errors calculated by bootstrap resampling of 25 independent SLiM2 simulations. The horizontal dashed line indicates α = −0.38, the best-fit α across the 25 UK Biobank traits.

## Discussion

We have quantified the MAF-dependent architectures of 25 diseases and complex traits under the 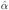 model^14, 15^ (Equation 1). We inferred negative values of α for all 25 traits and significantly negative values for 20 traits, corresponding to higher trait effects for lower-frequency SNPs. The best-fit distribution of α across traits had mean −0.38 (s.e. 0.02) and standard deviation 0.08 (s.e. 0.03), implying that only 8.9% (s.d. 2.7%) of SNP-heritability is explained by rare SNPs (MAF < 1%), despite significantly larger effects for rare variants. Although rare variants explain relatively little heritability, rare variant association studies may still identify variants of large effect that reveal interesting biology and actionable drug targets^11, 26^. On the other hand, rare variants will likely play only a limited role in polygenic risk prediction, which will be largely driven by common variants.

Using evolutionary modeling and simulations, we determined that the α model provides a good fit for common SNPs (MAF ≥ 5%), though it may potentially overestimate effects of rare SNPs; our estimate of 8.9% (s.d. 2.7%) of SNP-heritability explained by rare SNPs should therefore be viewed as an upper bound. We concluded that an average genome-wide negative selection coefficient on the order of 10^−4^ or stronger is required to explain the MAF-dependent architectures that we inferred. The best-fit α estimate across 25 traits implies an Eyre-Walker^9^ τ parameter between 0.3 and 0.5, quantifying the coupling between fitness effects and trait effects. Our finding that estimates of α (and hence τ) vary only modestly across traits is consistent with the action of pleiotropic selection, in which SNPs that affect the target trait also affect other selected traits^27, 28^; under direct selection, greater variation in τ would be expected, and traits that are not directly selected would have τ = 0.

Recent studies have investigated MAF-dependent architectures in genome-wide analyses of schizophrenia^3, 5^, as well as height and BMI^4^. These studies analyzed a small number of traits, and either did not analyze rare variants^3, 5^ or aggregated all MAF < 10% variants into a single MAF bin^4^, underscoring the difficulty of obtaining precise estimates of rare variant heritability using the MAF bin approach. Another study used targeted sequencing of 63 prostate cancer risk regions to conclude that 42% (s.e. 11%) of the prostate cancer SNP-heritability attributable to these regions in African Americans is due to rare SNPs (MAF < 1%), although rare variant heritability in Europeans was non-significant^6^.

A more recent study introduced a revised LDAK method^19^ (revising an earlier LDAK method^14^) and estimated a parameter that it referred to as α. We refer to this parameter as α_LDAK_, because it is different from the parameter α that was previously described in ref.^14, 15^ and that is defined and estimated in this paper. Specifically, the Discussion section of ref.^19^ states that the SNP effect size variance is proportional to [*p_j_*(1−*p_j_*)]^α^_LDAK_. However, that statement is incorrect. Actually, under the model of ref.^19^, the SNP effect size variance is proportional to [2*p_j_*(1−*p_j_*)]^α^_LDAK_ ·*w_j_*, where *w_j_* is an LD-dependent weight (see Equation 1 of ref.^19^). Unlike the LD-dependent weights that we use^17^, *w_j_* is dependent on MAF, with lower frequency SNPs having higher values of *w_j_*. Thus, SNP effect size is specifically not proportional to [*p_j_*(1−*p_j_*)]^α^_LDAK_, and α_LDAK_ is a parameter that is different from α. Indeed, our simulations confirmed that estimates of α_LDAK_ obtained using the LDAK software were upward biased by roughly 0.4 compared to the true α as defined in previous work^14, 15^ and this paper (see Supplementary Table 7). Thus, the revised LDAK method and software^19^ cannot be used to estimate α.

An unpublished study conducted in parallel to this work investigated MAF-dependent architectures of 28 UK Biobank traits^29^ using a Bayesian method to estimate a parameter identical to the α parameter that we estimate. Results of ref.^29^ were broadly similar to our results, but we note three key differences between the studies. First, ref.^29^ did not include rare variants (MAF < 1%) in their analyses, although we determined here that inclusion or exclusion of rare variants does not significantly affect our results. Second, ref.^29^ used an elegant approach to infer the polygenicity of each trait. Third, although ref.^29^ performed forward simulations to show that their findings implicate negative selection on trait-affecting SNPs, they did not use these simulation results to investigate the validity of their parametric inference model or to infer evolutionary parameters.

In addition, several recent studies have drawn inferences about evolutionary parameters that affect complex traits. Ref.^12^ and ref.^13^ estimated τ in type 2 diabetes to be approximately 0.1, by comparing the number of rare and low-frequency associations in empirical studies to the number in simulations. Ref.^6^ estimated τ by matching the heritability explained by rare SNPs (MAF < 1%) in their analysis of prostate cancer to simulation results, inferring 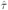 = 0.48 (95% CI: [0.19, 0.78]). We are not aware of any previous study that has drawn inferences about the genome-wide average strength of negative selection, although ref.^28^ used a different modeling approach to estimate the mutational target load.

We note several limitations in our work. First, our analyses are restricted to high-prevalence diseases and quantitative traits, as low-prevalence diseases are not well-represented in the UK Biobank due to random ascertainment. This motivates additional analyses of low-prevalence diseases, which could potentially be subject to stronger direct selection. However, we caution that our method might be susceptible to biases when used to analyze ascertained case-control traits, as previously described for linear mixed model based heritability estimation methods^30, 31^, meriting further investigation. Second, we use the Eyre-Walker model^9^ to parameterize the coupling between fitness effects and trait effects. The Eyre-Walker model has previously proven useful in a variety of settings^6, 12, 13^, but other coupling models are also possible^28, 32^. One limitation of the Eyre-Walker model is that it does not allow for signed correlations between SNP trait effect and selection coefficient, i.e. the damaging allele is equally likely to reduce or increase the trait value. This assumption is violated when the target trait is under direct selection, but is plausible if selection on the SNP is mainly pleiotropic, which appears to be the dominant form of selection for the traits analyzed here (see above). Third, we assume that the distribution of selection coefficients follows a gamma distribution. This assumption implies that there are no outlier SNPs under exceptionally strong negative selection. Such extremely selected SNPs would stay at very low frequencies and only affect our results if they had extreme effects on the target trait. However, such SNPs have not been identified for most complex traits^2^. We specifically assume that the distribution of selection coefficients has a gamma shape parameter of κ = 0.25, which is the value that ref.^22^ inferred for coding variants. Although we also considered different values of κ within the plausible range inferred by ref.^22^, it is possible that this parameter could be different for noncoding variants. However, we are not aware of any specific reason why this should be the case. Fourth, our analytic derivations ignore LD and assume a constant population size. Our derivations imply that α ≈ −2τ (see Online Methods), but our forward simulations, which include realistic LD patterns and demography, suggest that α ≈ −τ. The direction of this change is consistent with the action of background selection due to LD, since strong LD leads to a SNP’s frequency being influenced not only by its own selection coefficient but also by the selection coefficients of many other correlated SNPs, leading to a less negative α value for a given τ. However, this difference could potentially also be due to demography.

The impact of LD and demography on α could potentially be investigated further using forward simulations. Finally, our forward simulations assume that negative (purifying) selection is the dominant mode of selection affecting complex traits. Although positive selection is likely to affect some loci, recent work has suggested that selective sweeps were rare in human evolution^33^ and hence unlikely to have substantial genome-wide effects on MAF-dependent trait architectures. We also did not investigate the potential effects of stabilizing selection^28^. Despite these limitations, our quantification of MAF-dependent effect sizes and the underlying evolutionary parameters is broadly informative for the genetic architectures of diseases and complex traits.

## URLs

Software implementing our method will be released prior to publication as a publicly available, open-source software package at https://www.hsph.harvard.edu/alkes-price/software/; UK Biobank website, http://www.ukbiobank.ac.uk/; BGEN file format, http://www.well.ox.ac.uk/~gav/bgen_format/; UK Biobank genotype imputation manual, http://www.ukbiobank.ac.uk/wp-content/uploads/2014/04/imputation_documentation_May2015.pdf

## Acknowledgements

We are grateful to Ivana Cvijović, Kevin Galinsky, Alexander Gusev, Benjamin Neale and Nick Patterson for helpful discussions. This research was funded by NIH grants R01 MH101244 and U01 HG009088 and by a Boehringer Ingelheim Fonds fellowship. This research was conducted using the UK Biobank Resource under Application Number 16549.

## Online Methods

### Inferring frequency dependence of SNP effects

We assume a linear complex trait model for *N* individuals and *M* SNPs with

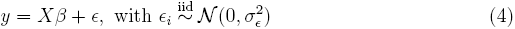

Here, *y* is a vector of *N* phenotype values with mean zero, *X* is the mean-centered genotype matrix, β is the vector of *M* SNP effects and ε is a vector of environmental effects (i.e. any non-SNP effects). Furthermore we assume the effect size of SNP *j* to be a random variable that follows a distribution depending on its minor allele frequency (MAF) *p_j_*:

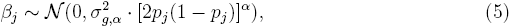

where effect sizes of two SNPs are independent conditional on their allele frequencies. A negative α value indicates larger trait effects on average for lower-frequency SNPs, whereas 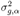 is the component of the SNP effect variance independent of frequency. This model, which we call the α model, has been used in previous analyses of complex traits^14, 15^. We note that β defines the per-allele SNP effect which is distinct from the heritability explained by a SNP. Under Hardy-Weinberg equilibrium and given Equation 5, the average heritability explained by a SNP of frequency *p* is proportional to [2*p*(1 − *p*)]^1+α^.

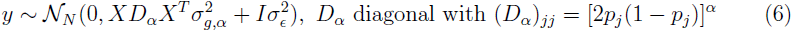

Given the genotype matrix *X*, SNP frequency vector *p* and phenotype vector *y*, the likelihood over the three parameters 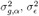 and α is fully defined by Equation 6. Hence, the MLE of the parameter triple (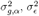, α) can be found directly by maximizing the corresponding likelihood. Since we are primarily interested in estimating α, we used a profile likelihood based approach, with the profile likelihood of α defined as 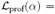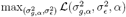. In this analysis we use 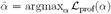 as the estimator of α, given genotype and phenotype data *X* and *y*. 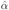 is also equal to the α value in (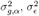, α) that maximizes the total likelihood in Equation 6.

In practice, the profile likelihood 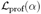 was derived in the following way: for some 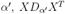, 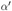 was calculated. Given phenotype values *y* and for a given 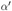, we inferred maximum likelihood estimates for 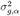 and 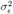 via restricted maximum likelihood estimation^34^, using the GCTA software implementation^35^. This procedure was repeated for a range of 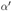. Here we used a minimal range of 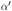 ∈ {−1.00,−0.95,…, 0.00} for all traits, but extended the range to higher values if necessary, such that there is a minimal difference of 5 in log profile likelihood between the mode and the boundary. This ensures that the part of the curve that is significantly above zero is sampled. These data points were then interpolated with a natural cubic spline, yielding the final profile likelihood curve. Credible intervals for 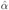 were estimated by combining the profile likelihood curve with a flat prior. Although our above modeling assumes a quantitative trait, this method is equally applicable to randomly ascertained case-control traits since all likelihood calculations are performed using the GCTA software, which analyzes case-control traits accordingly via a liability threshold model^36^.

Given 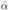 for a set of phenotypes, the cross-trait estimate, 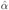_cross-trait_, was calculated as the inverse standard error weighted mean across the traits. We tested for heterogeneity of true underlying α values across n traits by comparing 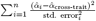 to a 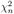 null statistic. The best-fit standard deviation in true α values across traits, was calculated by assuming normally distributed true α with mean 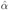_cross-trait_, and then choosing the standard deviation, for which the variance of the simulated 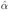 using the inferred standard errors matched the variance of the 25 α estimates most closely.

### Correcting for LD-dependent architectures

Ref.^17^ showed that for a given MAF, SNPs with higher LD have lower per-allele effects on average. Specifically, they use level of LD (LLD), defined as the rank-based inverse normal transform of the LD score. LLD is transformed separately in each part of the MAF spectrum, ensuring that it is independent of MAF. Ref.^17^ reported that SNPs that have LLD one standard deviation above the mean have a squared per-allele effect size reduced by (30±2)% on average. This violates our assumption that, at a given MAF, all SNP effects are independent and identically distributed.

To avoid bias in our estimation due model misspecification, we incorporated LD-dependent SNP effects by changing Equation 1 to

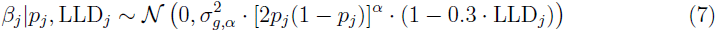

This expression incorporates the LD dependence of ref.^17^, however, since LLD has mean zero and is independent of MAF, 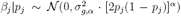 still holds, even though effect sizes β are not iid given *p*. To remove the LD dependence in the effect size distribution, we calculated a renormalized genotype matrix 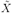, with 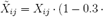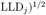. This effectively changes the complex trait model in Equation 4 to 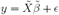, where now 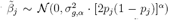 is again iid for a fixed *p*. Unless otherwise stated, we hence estimated α using 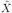 instead of *X* to avoid biases due to LD-dependent architectures.

### Genotype data

We use the UK Biobank phase 1 data release (see URLs), which comprises of data from 152,729 individuals genotyped at 847,131 SNP loci. Here, we only used data from 113,851 individuals following selection criteria previously used by Galinsky et al.^37^: individuals were selected to have self-reported and confirmed British ancestry and related individuals were removed from the analysis such that the pairwise genetic relatedness is < 5% (after LD-pruning SNPs). Individuals that had withdrawn consent to participate in the UK Biobank project after initial publication were removed from the analysis. We used imputed genotype data as provided by UK Biobank. These genotypes were imputed using the IMPUTE2 software^38^ and a joint reference panel from the UK10K project^39^ and 1000 Genomes Phase 3^40^. The resulting imputed genotype data includes roughly 70,000,000 SNPs loci across the 22 autosomal chromosomes. The data was downloaded in the BGEN file format (see URLs), a compressed file format that includes – for each individual and variant site – the probability of being homozygous reference, heterozygous, or homozygous alternative. Due to imputation uncertainty, the genotype matrix *X* and the allele frequencies *p* are not known precisely. Instead, we use the expected genotypes given these probabilities (genotype dosages). To exclude large-effect SNP loci from human leukocyte antigen genes, SNPs on chromosome 6 in the 30-31Mb region were masked and we verified that no significant associations were found in nearby regions after masking. Due to memory constraints, GCTA could not be run using a GRM of all 113,851 individuals at once. Instead, we divided all individuals into 3 equally sized batches, calculating the profile likelihood of α for each batch and using the sum of the resulting log likelihoods to compute the final likelihood curve.

Although our analysis does not require knowing all imputed genotypes precisely, we do assume that the genotype probabilities are well calibrated, i.e. that we are not overly confident in the imputation accuracy. Since imputation accuracy is difficult to assess if the number of minor alleles in the reference panel is very low, we only used SNP loci that had 5 or more minor alleles in the UK10K reference panel (MAF > 0.07%) in our main analysis. To further assess calibration of imputation noise, we compared the uncertainty implied by the genotype probabilities with an empirical assessment of imputation accuracy performed by the UK Biobank study (see URLs). Supplementary Figure 3 shows that imputation accuracy is significantly overestimated for SNPs of frequency 0.1% or less, which could potentially bias our results. However, repeating α estimation only using SNPs of MAF > 0.3% did not lead to significantly different results, implying that our results are not significantly affected (see Supplementary Table 3).

### Simulations

Simulations were performed using genotype data from an *N* = 5,000 random subset of the 113,851 unrelated British UK Biobank individuals. We used *M* = 100,000 consecutive SNPs from a 25Mb block of chromosome 1. *N* and *M* where chosen such that the simulations had similar statistical power as the main analysis^41^. As in the main analysis, only SNPs with at least 5 minor alleles (MAF > 0.07%) in the UK10K reference panel were included. Phenotype values were generated using the linear model described in Equation 4. The trait effect of the *j*^th^ SNP was drawn from 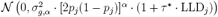, with τ* = −0.3 when simulating LD-dependent architectures^17^, and τ* = 0 otherwise. The environmental noise variance was chosen such that the simulated trait had the desired heritability. In simulations with only 1% of SNPs causal, the causal SNPs were chosen at random. Imputation noise was introduced by randomly sampling the genotypes used to simulate phenotypes from imputed genotype probabilities, as reported by UK Biobank. In simulations without imputation noise, genotype dosages, i.e. the expected number of minor alleles, were used. In the inference procedure, we used genotype dosages in both types of simulations.

Simulations to estimate α_LDAK_ were performed using the same set of 5,000 individuals and 100,000 SNP loci. Phenotype values were simulated as described above. α_LDAK_ estimation was performed in the same way as in the previous set of simulations, only now using the LDAK software^19^ to calculate the likelihood for a given α value instead of the GCTA software. This approach hence includes the LD weights proposed by LDAK and is identical to their proposed approach for estimating α, although, to enable a more accurate comparison, we used a finer set of tested α values (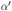 ∈ {−1.00, −0.95,…, 0.60}) than in their study (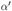 ∈ {−1.25, −1.00, −0.75, −0.50, −0.25, 0.00, 0.25}). Due to computational constraints we did not use their workflow for imputed genotypes, but rather used the same hard-called genotypes for both phenotype simulations and estimation, an option available in LDAK.

### Correcting for bias in heritability estimation

Heritability estimation methods based on standard restricted maximum likelihood (REML) estimation in a linear mixed model framework^16^ require that all SNP effects are iid distributed in order to avoid biases. In the case of MAF-dependent SNP effects, this assumption is clearly broken. This issue has been addressed in previous work and several solutions to this problem have been suggested^15, 19^. Here we show that knowing α for a given trait can provide another way to avoid heritability estimation biases due to MAF-dependent architectures. As previously stated, our model assumes 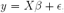, with 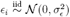 and 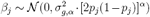. Here β is the per-allele effect, the average effect on the phenotype of having one minor allele. However, one can define renormalized genotypes 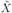, with 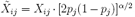. The per-normalized-allele effects are now 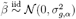 in 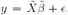. Since 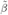 are now iid, 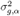 and 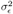 can now be estimated without bias from 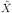 and *y* using REML. The variance in the phenotype explained by *M* SNPs can be calculated in the following way:

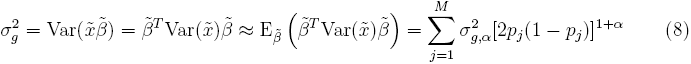

where 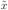 is a random renormalized genotype row vector. Here we used the fact that 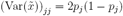 under Hardy-Weinberg equilibrium and cross terms cancel since 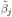 are independent and mean zero. We define 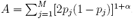, with the genetic variance 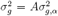. If α = −1, as has been used in many previous methods^14, 16^, *A* is simply equal to *M*.

In practice, heritability estimation was performed in the following way: the renormalized genotype matrix 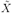 was calculated using the 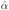 as estimated from the data. From 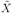 and the phenotype vector, 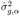 and 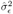 were obtained using GCTA REML^35^. Our SNP heritability estimate 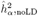 is then defined as 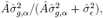, with 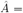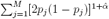. 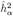 was calculated equivalently only now including previously described LD weights, i.e we used 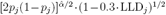 instead of 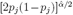 when calculating 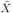 and *Â*.

### Phenotype selection and preprocessing

In this analysis we investigated 25 highly heritable and polygenic human traits (see Table 2) from the UK Biobank study (see URLs). Specifically, we required a SNP heritability of 0.2 or more for quantitative traits and 0.1 or more for case-control traits (on the observed scale, see ref.^36^), as well as at least 50% of the 113,851 British ancestry individuals to be phenotyped. We also removed phenotypes for which the top 10 SNPs explained 10% or more of the trait variance, so as to avoid α estimates that are dominated by a few top SNPs, as our goal is to study polygenic architectures. (Only one trait, mean platelet volume, was removed due to this restriction.) The 25 traits that we chose include 21 quantitative traits and 4 case-control traits. 11 of the quantitative traits are blood cell traits, whereas the remaining 14 include a wider range of physiological measurements and diseases. Since the number of available blood cell traits was large and many of them were highly correlated, we additionally required blood cell traits to have a pairwise phenotypic correlation of *r*^2^ < 0.5, removing the less heritable trait for any correlated pair.

For each trait, phenotype values had outliers removed and fixed effects were regressed out. Specifically, phenotype values 4 or more standard deviations away from the mean (or similarly extreme outliers for skewed distributions) were removed from the analysis. Sex and 10 principal components of the GRM were included as fixed effects for all traits, with additional trait specific covariates also included for some traits (see Supplementary Table 8). All trait values were then rank-based inverse normal transformed before being analyzed.

### Inference of fitness-trait coupling and selection parameters

We aimed to use the frequency dependence of SNP effects to draw conclusions about the fitness effects of SNPs, as well as the coupling between between fitness and the target trait effects. Let β^2^|p be the squared trait effect size of a SNP given its MAF *p*, and *s* the fitness effect of the SNP, which is here assumed to be deleterious or neutral. From the law of total expectation it follows that E(β^2^|*p*) = E (E (β^2^|*s*, *p*) |*p*). The main assumption of this analysis is that, at a given selection coefficient, the effect size of the SNP is independent of its frequency, i.e. E(β^2^|*s*, *p*) = E(β^2^|*s*). This is equivalent to the statement that the frequency dynamics of a SNP is inuenced by β^2^ only through *s*. We then use the model of Eyre-Walker^9^, where the absolute value of β is proportional to *s^τ^* (1+∈), with 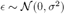 and τ indicating how strongly β depends on *s*. It follows that E(β^2^|*s*) ∝ *s*^2τ^ and from above, for some constant *c*,

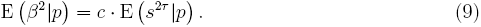

Given a positive τ, this equation shows that increased average effects of lower-frequency SNPs requires lower-frequency SNPs having increased *s* and hence implies significant negative selection. Some previous analyses^4, 6, 29^ have argued that in the absence of selection, SNPs of MAF ranges of equal width (e.g. 5-10% and 10-15%) are expected to explain an equal fraction of heritability. However, even in the absence of selection, population expansion can lead to excess rare variants, leading to increased rare variant heritability^42^. Increased rare variant heritability is therefore not necessarily a sign of selection.

Assuming we know τ and the joint distribution of *s* and *p*, E (β^2^|*p*) can be derived from Equation 9. We simulated samples of this distribution using the evolutionary forward simulation framework SLiM2 (ref.^23^). Simulations were run with a European demographic model inferred by ref.^24^, a burn-in of 3,880 generations before the bottleneck, a mutation rate of 2 · 10^−8^ per base pair per individual per generation^43^, and a recombination rate of 10^−8^ per base pair per individual per generation^44^. These simulations also require assumptions about the distribution of fitness effects (DFE), i.e. the distribution of *s* for de novo mutations, but the DFE for genome-wide SNPs in humans is currently not known. We assumed a gamma distributed DFE, using a plausible range of average fitness effects, 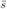 ∈ {10^−3^, 10^−4^, 10^−5^}, and shape parameters of 0.125 and 0.25 which includes the range of plausible values derived by ref.^22^. For each choice of DFE we simulated 25 independent replicates over a 4Mb block each, for a total of 100Mb with each DFE. In all simulations the Eyre-Walker noise parameter, σ^2^, was set to zero. This parameter does not change SNP effects on average and is therefore negligible in the limit of large SNP numbers. This was also noted in original analysis by ref.^9^.

In the absence of LD between selected SNPs and assuming a constant effective population size Ne, E(β^2^|*p*) can also be derived analytically. Under these assumptions and assuming mutation rate per base pair μ « 1/*N_e_* (ref.^43^), it is known that P(*p|s*) ∝ [*p*(1−*p*)]^−1^e^−4*N_e_sp*^ (ref.^45^). Given *s* is drawn from a gamma distribution with mean 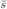 and shape parameter κ, we obtain

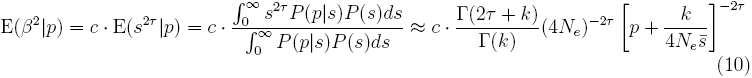

This result shows that for 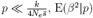, E(β^2^|*p*) is constant, whereas for 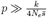 it falls off as *p*^−2τ^. We note that these calculations imply α ≈ −2τ, whereas α is significantly less negative in simulations (see Figure 3), with the difference likely being due to LD between SNPs with different selection coefficients (see Discussion). For simplicity, we have here assumed that p is the derived allele frequency – if *p* is the minor allele frequency, results are similar though there is a correction factor for very common SNPs, roughly matching the (1 − *p*) factor in the our E(β^2^|*p*) ∝ [*p*(1 − *p*)]^α^ model (see Supplementary Figure 5).

When fitting α to SNP effects from a simulation with a given 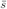, κ and τ in Figure 3, we only used SNPs with frequency above 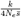. 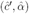 is calculated by minimizing the squared deviation between 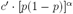 and the simulated SNP effects summed over all SNPs from 25 independent simulations. Error bars were obtained by bootstrap resampling of these 25 simulations. The proportionality constant in Equation 9 does not affect 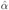 and was set to *c* = 1. When estimating τ from 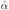 of a given trait, we assumed a flat prior on α over [−1,0] and on τ over [0,1], in which case 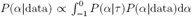. Here, P(α|data) is proportional to the calculated profile likelihood and P(α|τ) is based on estimates and error bars displayed in Figure 3, assuming equal probability for 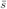 = 10^−3^ and 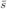 = 10^−4^, and κ = 0.25. Using κ = 0.125 lead to similar results, e.g. α = −0.38 then corresponds to τ ∈ [0.33,0.43] instead of τ ∈ [0.32,0.48] for κ = 0.25.

